# Hypoxia-Induced PIM Kinase and Laminin-Activated Integrin α6 Mediate Resistance to PI3K Inhibitors in Bone-Metastatic CRPC

**DOI:** 10.1101/685602

**Authors:** Rachel K. Toth, Jack D. Tran, Michelle T. Muldong, Eric A. Nollet, Veronique V. Schulz, Corbin Jensen, Lori A. Hazelhurst, Eva Corey, Donald Durden, Christina Jamieson, Cindy K. Miranti, Noel A. Warfel

## Abstract

Bone-metastatic castration-resistant prostate cancer (CRPC) is lethal due to inherent resistance to androgen deprivation therapy, chemotherapy, and targeted therapies. Despite the fact that a majority of CRPC patients (approximately 70%) harbor a constitutively active PI3K survival pathway, targeting the PI3K/mTOR pathway has failed to increase overall survival in clinical trials. Here, we identified two separate and independent survival pathways induced by the bone tumor microenvironment that are hyperactivated in CRPC and confer resistance to PI3K inhibitors. The first pathway involves integrin α6β1-mediated adhesion to laminin and the second involves hypoxia-induced expression of PIM kinases. *In vitro* and *in vivo* models demonstrate that these pathways transduce parallel but independent signals that promote survival by reducing oxidative stress and preventing cell death. We further demonstrate that both pathways drive resistance to PI3K inhibitors in PTEN-negative tumors. These results provide preclinical evidence that combined inhibition of integrin α6β1 and PIM kinase in CRPC is required to overcome tumor microenvironment-mediated resistance to PI3K inhibitors in PTEN-negative tumors within the hypoxic and laminin-rich bone microenvironment.

## INTRODUCTION

Disseminated prostate cancer is ultimately lethal, because these metastases almost always display inherent or acquired resistance to current therapies. Despite recent improvements in androgen deprivation therapy (ADT), e.g., enzalutamide and abiraterone, for the treatment of advanced prostate cancer, almost all patients ultimately develop castration-resistant disease (CRPC). Moreover, CRPC is not only resistant to ADT, but also to chemotherapy and targeted therapies. Since ~70% of CRPC patients harbor a constitutively active PI3K survival pathway, primarily due to PTEN loss [1], it was highly anticipated that specific targeting of PI3K/mTOR would be an effective strategy to treat CRPC. However, despite the dependency of prostate cancer on PI3K signaling for survival, fewer than 25% of CRPC patients displayed progression-free survival, and no objective responses were observed with PI3K or mTOR inhibitors [1]. Furthermore, clinical trials combining ADT and PI3K inhibitors in CRPC produced limited responses [1]. Resistance to PI3K inhibitors is likely due to the presence of constitutively active androgen receptor (AR)-driven survival pathways that are no longer inhibited by androgen blockade in CRPC. Inhibitors of several AR-independent survival pathways, including receptor tyrosine kinases, src kinases, and survivin, have been tested clinically [2–5], but none were able to overcome CRPC as single agents, with or without ADT, achieving <25% stable disease and no cures. In these trials, patients were not stratified with respect to elevated expression of the specific targets, and no analyses were performed to determine the extent of target inhibition. Therefore, a fundamental concept is lacking in our understanding of the survival signaling mechanisms that operate in CRPC, and our current strategies are not sufficient to determine whether targeted therapies will be effective.

Prostate cancer commonly metastasizes to the bone, accounting for over 80% of all disseminated CRPC. The bone tumor microenvironment (TME) is vastly different than the site of primary tumor. Two bone-enriched environmental factors, the extracellular matrix (ECM) and hypoxia, are independently established mechanisms for drug resistance, but their mechanistic role in CRPC or drug resistance in bone metastasis in general has received limited attention. Cell adhesion to the ECM by integrins is a major mediator of chemotherapy and radiation resistance in many cancers, including prostate cancer [6–8]. Blocking adhesion to the ECM sensitizes prostate cell lines to both radiation and chemo therapies [9–11]. Several mechanisms have been proposed for how cell adhesion-mediates drug resistance (CAM-DR), including integrin-mediated activation of JNK, kindlin-2, FAK, Stat3, and MDR1 [12–17]. CAM-DR is also a resistance mechanism for targeted therapies. Upregulation of integrin α2β1 confers resistance to mTOR inhibitors [18], which could explain the failure of mTOR inhibitors in clinical trials for CRPC. Integrin ligands and FAK activation were shown to confer resistance to cabozantinib in metastatic prostate cancer [19]. Thus, integrin-mediated adhesion activates multiple mechanisms to promote drug resistance.

Another important mediator of drug resistance within the bone TME is hypoxia [20]. Hypoxia is especially relevant to prostate cancer, as the prostate gland is hypoxic compared to many other soft tissues, and hypoxia increases with stage in prostate cancer [21, 22]. Furthermore, the bone TME is intrinsically hypoxic. Thus, hypoxia provides a robust mechanism for escaping ADT and promoting CRPC. Attempts to therapeutically target hypoxic tumors have focused largely on inhibiting HIF-1. These drugs are in clinical trials for prostate cancer, but to date none as single agents have improved overall survival [23]. PIM kinases are a family of oncogenic Ser/Thr kinases whose levels are elevated in high-grade prostatic intraepithelial neoplasia relative to normal tissue and are further elevated in CRPC [24, 25]. It was recently reported that hypoxia enhances the expression of PIM1 and PIM2 in prostate cancer cells, independent of HIF-1 and *PIM* mRNA transcription [26]. Moreover, small molecule inhibitors of PIM selectively kill hypoxic prostate cancer cells *in vitro* and *in vivo* by increasing reactive oxygen species (ROS) to toxic levels [26]. Thus, activation of PIM kinases represents a HIF-1-independent pathway that is critical for cell survival during hypoxia [26, 27]. Moreover, PIM expression has been described as a drug resistance mechanism for PI3K inhibitors in prostate and breast cancer [28, 29].

In this study, we identify two survival pathways induced in the hypoxic and laminin-rich bone TME that are hyperactivated in CRPC and confer resistance to PI3K inhibitors. The first pathway involves integrin α6β1 and adhesion to laminin, which prevents cell death and lowers oxidative stress. The second pathway involves hypoxia-induced, but HIF-1-independent, expression of PIM kinases. We show that both pathways are selectively activated in metastatic CRPC patient-derived xenograft (PDX) models. Since each pathway prevents cell death and reduces oxidative stress, they appear to work in parallel. We further demonstrate that each of these pathways drives resistance to PI3K inhibitors in PTEN-negative models of CRPC. Thus, combined targeting of these independent TME-driven survival pathways in combination with PI3K represents a promising approach to treat PTEN-negative bone-resident CRPC.

## MATERIAL AND METHODS

### Cell lines and tissue samples

PC3 and LNCaP cell lines were purchased from ATCC (Manassas, VA, USA). The C4-2 cell line was obtained from Robert Sikes, University of Delaware [30]. All cell lines were validated yearly by Single Tandem Repeat sequencing. Cell lines were maintained in RPMI supplemented with 10% FBS, 1 mM sodium pyruvate, 2 mM glutamine, 0.3% glucose, 10 mM HEPES, and 30 U/mL Pen/Strep. The human PDX PCSD1 was derived from a patient femoral metastasis and established as previously described [31, 32]. Human PDX tumors and fixed samples from LuCaP 23.1, LuCaP 23.1CR, LuCaP 35, LuCaP 35CR, LuCaP 77, LuCaP 77CR, LuCaP 86.2, LuCaP 105, and LuCaP 105CR have been previously described [33].

### Chemicals and reagents

MTI-101 was supplied by Modulation Therapeutics (Morgantown, WV, USA), SF1126 by Signal Rx (Omaha, NE, USA), and PX866 by Oncothyreon (Bothell, WA, USA). Laminin was purchased from Corning (Corning, NY, USA), R1881 and Enzalutamide (MDV3100) were purchased from Sigma (St. Louis, MO, USA), rat polyclonal GoH3 integrin α6 blocking antibody was purchased from BD Pharmingen (San Jose, CA, USA), and AZD1208 was purchased from AdooQ Bioscience (Irvine, CA, USA).

### Antibodies

The following antibodies were used for western blotting: HIF-1α, tubulin, and actin were purchased from BD Transduction Laboratories. PIM1 (D8D7Y), PIM2 (D1D2), PIM3 (D17C9), and p-Akt (S473) antibodies were purchased from Cell Signaling Technologies (Danvers, MA, USA). The antibody against Nrf2 (sc-13032) and integrin β1 (sc-374429) were purchased from Santa Cruz Biotechnology (Dallas, TX, USA). Integrin α6 rabbit polyclonal antibody (AA6A) was obtained from Dr. Anne Cress, University of Arizona [34]. The following antibodies were used for immunohistochemistry (IHC): HIF-1α (nb100-105; Novus Biologicals; Centennial, CO, USA), PIM1 (ab75776; Abcam; Cambridge, UK), BNIP3 (EPR4034; Abcam), pAkt (S473) (Cell Signaling Technologies), and rabbit polyclonal anti-integrin α6 (AA6NT) from Dr. Anne Cress [35].

### In vivo mouse studies

Male SCID and NSG mice were housed and maintained in accordance with the Institutional Animal Care and Use Committee and state and federal guidelines for the humane treatment and care of laboratory animals. For subcutaneous cell line xenograft models, PC3 cells were injected into the rear flanks of NSG mice at a density of 1 × 10^6^ cells per injection in PBS in 100 μL total volume. For bone-resident tumors, 1 × 10^6^ cells were resuspended in 10 μL of PBS and injected into the femur or tibia of 5-to 6-week-old male NSG mice. As a control, 10 μL of PBS was injected into the contralateral femur. In both cases, tumors were harvested 4-6 weeks after injection and processed for immunohistochemical staining.

For PDX models, including LuCaP 23.1, LuCaP 35, LuCaP 77, LuCaP 86.2, and LuCaP 105, as well as the castration-resistant derivatives, tumors were maintained by transplantation of tumor fragments under the skin or injection of single cell suspension into the tibia of SCID mice, as previously described [33]. Isolated tumor tissues from subcutaneous or tibial injections were processed for IHC. For PDX tumor drug treatments, subcutaneous tumors were grown to 100 mm^3^ and mice were randomly assigned to two or four treatment groups, with 14 mice per group: vehicle (10% DMSO in PBS), PX866 (2 mg/kg p.o. 3x/wk), ITGα6 Ab (10 µg GoH3 anti-integrin α6 antibody i.p. 1x/wk), or both combined. In addition, PX866 was combined with MTI101 (40 mg/kg p.o. 3x/wk). All groups were treated for 28 days, and tumor volumes were monitored twice a week by caliper measurement. At the end of the study, tumors were fixed, embedded in paraffin, and processed for IHC staining.

### PCSD1 PDX model

All studies with human samples were performed with the approval of the University of California, San Diego School of Medicine Institutional Review Board. The patient provided written informed consent. The PCSD1 PDX model was generated as previously described from prostate cancer bone metastasis cells that were isolated directly from the bone metastatic lesion in the proximal femur of a patient with CRPC having a hemi-arthroplasty to treat a pathologic fracture in the right femur head [31, 32]. The Gleason score of the original tumor was 10 (5+5), and mixed osteoblastic and osteolytic lesions were observed in the proximal femur on computed tomography in the patient. For experiments here, PCSD1 tumor cells were freshly isolated from low passage intra-femoral (IF) xenograft tumors in NSG mice and single cell suspensions prepared as previously described [31, 32]. Briefly, the IF xenograft tumor specimen was minced by surgical scalpel into small pieces and disaggregated by digestion in Accumax (Millipore) then filtered through a sterile 70 µm mesh filter. Dissociated cells were centrifuged at 1200 RPM, 5 minutes at 4°C, washed three times and resuspended in Iscove’s modified DMEM media with 10% FBS. Cells were mixed 1:1 with high concentration BD Matrigel Basement membrane Matrix (BD Biosciences) for IF injection of 50,000 cells in 15 µL.

### Direct intra-femoral injection in mice

Male, 6-8-week old NSG mice purchased from Jackson Laboratories (NOD.Cg-Prkdcscid Il2rgtm1Wjl/SzJ, Jackson Labs Stock # 005557 (NSG)) were used for direct IF injection. Mice were anesthetized with intra-peritoneal (i.p.) injection of a mix of 100 mg/kg ketamine and 10 mg/kg xylazine. The right knee was held in flexed position and a 25-gauge needle (Monoject 200 25 × 5/8A) was inserted into the femoral condyle until there was no resistance. It was used as a porthole for injection of 15 µL sample using a 0.3 mL syringe and 27-gauge needle. Mice were monitored weekly for health, body weight, and appearance of palpable tumor and the length and width of tumors were measured with calipers and volume measured in by in vivo bioluminescence imaging system (IVIS 200; Caliper Inc.) [31]. For treatments, tumors were grown for 5 weeks and mice randomly assigned to treatment groups, 10 mice per group. Mice were treated with vehicle, enzalutamide (o.g. 50 mg/kg/day), or SF1126 (o.g. 50 mg/kg 3x/wk) for 4 weeks and tumor growth measured by caliper or IVIS. Mice were sacrificed at 28 days of treatment or when tumor length reached 1.5 cm, the maximal allowable size according to UCSD ACP standards.

### Micro-computed tomography: image acquisition, processing and analysis

Mice were sacrificed 8-10 weeks after IF tumor cell injection. All samples were cut down under the lumber vertebra and fixed in 10% neutral buffered formalin. Micro-CT images were acquired using the µCT SkyScan 1076 (Bruker, Kontich, Belgium) at a source power of 59kV/167µA and spatial resolution of 9.06 µm/pixel with a 0.5-mm-thick aluminum filter. The rotation was set at 0.7 degrees per step for 180 degrees. Image processing: The reconstructions were performed with NRecon software package (SkyScan, Bruker, Kontich, Belgium) to obtain transaxial grayscale images. Each region of interest (ROI) was delineated using CT-Analyzer software (SkyScan, Bruker) so as to include both sides of the whole femur. Two-dimensional (2D) transaxial, coronal, and sagittal images were obtained with Data Viewer software (SkyScan, Bruker) based on the ROI data.

### Immunohistochemistry

Formalin fixed samples were deparaffinized with xylene/ethanol then antigen retrieval performed using Dako S1699 Retrieval Buffer or 1 mM EDTA in water at pH 8.0 at 90°C for 30 mins. After a 5 min peroxidase blockade, slides were blocked for 10 min at R.T. using filtered 0.5% BSA in PBS, filtered 5% goat serum in PBS, 1X mouse FcR blocker (stock: 50x Mac’s Miltenyi Biotec 130-092-575), and 1X human FcR blocker (stock: 50x Mac’s Miltenyi Biotec 130-059-901). After washing, slides were incubated with primary antibody overnight at 4°C. IHC DAB staining kit (TL-015-HD by Invitrogen) was used for final detection, counterstained with hematoxylin, and mounted in non-aqueous solution (Richard-Allen Scientific 4112).

### Analysis of immunohistochemistry

Tissue images underwent ImageJ/Fiji color deconvolution to separate the blue and brown pixel channels, representing DAPI and the protein of interest, respectively, from the raw images. Specifically, we utilized the IHC Profiler plugin [36] with the mode set to “Cytoplasmic Stained Image” and using the “H DAB” vector. Once deconvolution was complete, the H DAB channel images were further analyzed using the IHC Profiler Macro to assess the percentage of pixels contributing to negative, low positive, positive, and high positive staining. The values for positive and high positive were combined to determine a new value for percentage of pixels positive for our given staining conditions.

### In vitro assays

For assays done on laminin, plates were precoated with 10 µg/mL at 37°C for 1 hour and then treated with 5% BSA prior to plating the cells. For R1881 treatment, cells were serum-starved in media containing 0.1% charcoal-stripped serum for 24 hours prior to treatment. When indicated, cells were maintained in a hypoxic environment (1.0% O_2_) using a hypoxia workstation (In vivo2 400; Ruskinn; Sanford ME, USA). Immunoblotting was performed as described previously [37]. For ROS assays, C4-2 cells were plated in 6-well plates and cultured in normoxia or hypoxia ± AZD1208, SF1126, or MTI-101 for the indicated times. Then, the cells were washed with PBS and incubated with CM-H2DCFDA (1 μmol/L in PBS; Molecular Probes; Eugene, OR, USA) for 10 minutes. The CM-H2DCFDA was removed and the cells were washed once with PBS. The cells were immediately harvested by trypsinization, centrifuged, and resuspended in PBS for FACS analysis (FACSCanto II; Beckon Dickinson; Franklin Lakes, NJ, USA). To measure viability, the indicated cells were plated in 96-well plates and treated with AZD1208, MTI-101, or SF1126 at their respective IC_50_ or the indicated combinations in normal (20% O_2_ on plastic) or bone-enriched environmental (1.0% O_2_ and laminin) conditions. After 48 h incubation, living cells were stained with crystal violet or trypan blue, according to previously described methods [38].

### Statistical methods

All Western blots shown are representative of at least three independent experiments. Differences between independent groups were determined by the Student’s *t* test and linear regression analysis. Two-way ANOVA was used to analyze differences in survival between groups with two independent variables (i.e., normoxia vs. hypoxia). For *in vivo* experiments, univariate cross-sectional analyses compared tumor volume by treatment group using the Wilcoxon Rank Sum rank test. After cube root transformations to achieve approximate normality, multivariate analysis compared tumor volume by day of observation, measurement time, and treatment using mixed effects models that accounted for the correlations between the measurements obtained over time for individual mice. All analyses used STATA 15 (StatCorp, College Station, TX, USA). All data are presented as the mean ± standard error, and a *p* < 0.05 was considered to be statistically significant.

## RESULTS

### The bone TME is hypoxic and promotes resistance to PI3K inhibitors

Clinical data has shown that PI3K inhibitors do not significantly increase patient survival in patients with constitutive activation of the PI3K pathway due to PTEN loss or other genetic alterations, suggesting that TME-driven signaling may play a role in promoting therapeutic resistance in bone-resident CRPC. Hypoxia is an established mechanism for drug resistance in solid tumors, but the extent and importance of hypoxia in bone-metastatic disease has not been well described. To determine the relative contribution of hypoxia in bone-resident tumors vs. subcutaneous models, PC3 cells, LuCaP 35CR, and LuCaP 86.2 PDX tumors were implanted subcutaneously (SC) or into the tibia or femur (Bone) and allowed to grow for 4 weeks. Sections from SC and Bone tumors were stained for HIF-1α, an established marker of hypoxia (Fig. 1A). The relative extent of hypoxia was calculated as the percent HIF-1α-positive area within each tumor sample. HIF-1α staining was significantly higher in bone-resident tumors (Fig. 1B), indicating that tumors growing in the bone TME are exposed to significantly more hypoxia than subcutaneous tumors.

**Figure 1.**
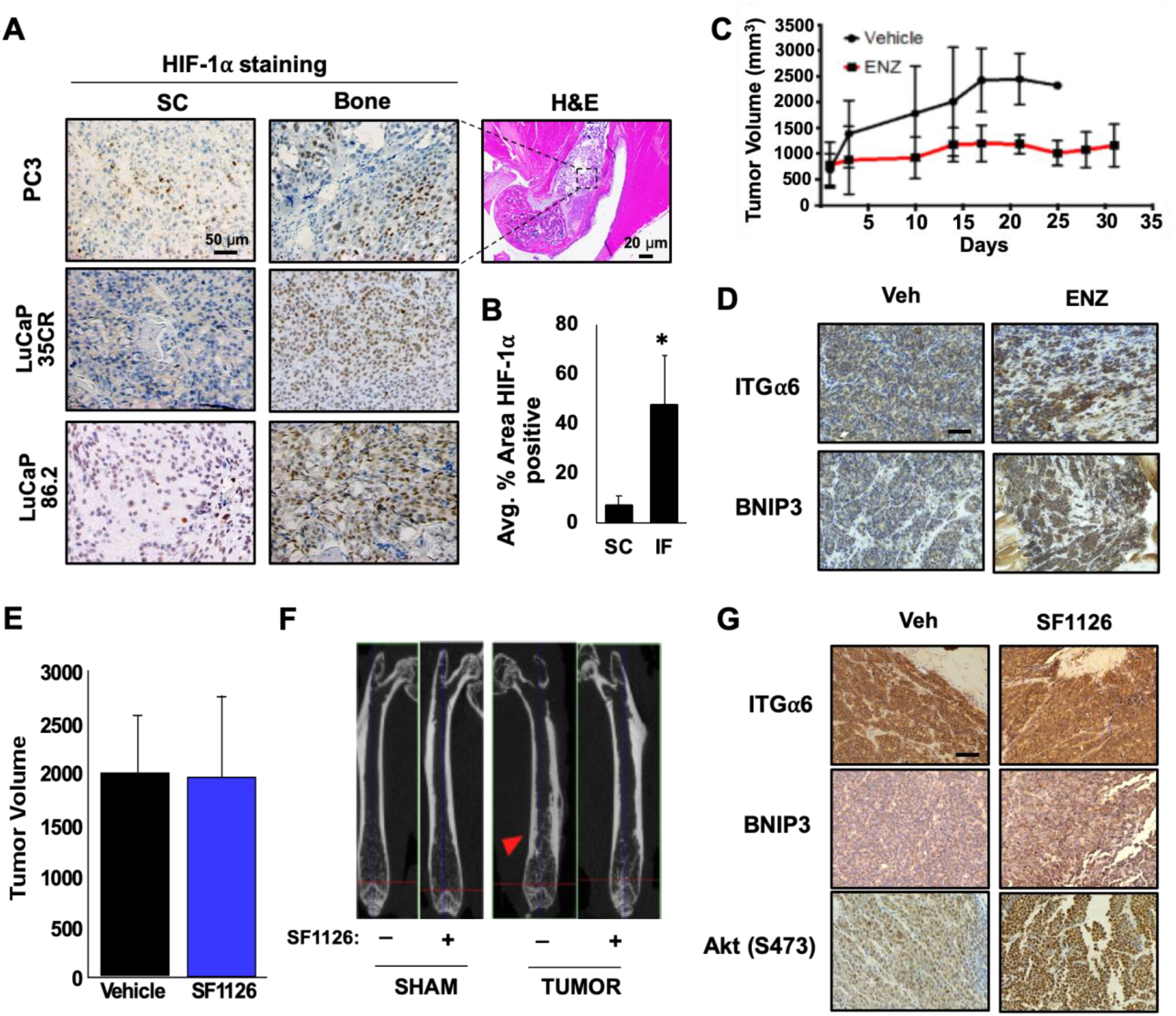
The bone TME is hypoxic and promotes resistance to PI3K inhibitors in CRPC. **A)** PC3 cells or two different LuCaP PDX tumors were implanted subcutaneously (SC) or in bone (Bone) and grown for 4 weeks. Tumors from each group were fixed and immunostained for HIF-1α, and **B)** positive staining was quantified. **C)** PDX PCSD1 was injected intrafemorally into NSG mice, and mice were treated for approximately 4 wks with vehicle or enzalutamide (ENZ; 50 mg/kg/day) and tumor growth measured by caliper. **D)** Tumors from each group were fixed and stained for ITGα6 and BNIP3. **E)** PDX PCSD1-injected mice were treated 5 weeks later with or without 50 mg/kg SF1126 3x/wk for 4 wks and growth assessed by IVIS. **F)** MicroCT imaging of bone from vehicle- or SF1126-treated mice. Red arrow; osteoblastic lesion. **G)** Tumors from each group were harvested, fixed and stained with the indicated antibodies. * *p* < 0.05; n=10; error bars indicate SEM.

To determine how bone affects resistance to androgen deprivation therapy (ADT) and PI3K inhibitors, we utilized the PCSD1 PDX tumor line, which was established from a PTEN-negative human bone metastasis biopsy and propagated in the femurs of NSG mice [32]. We previously demonstrated that the PCSD1 model accurately reflects the resistant phenotype observed in patients; bicalutamide completely suppresses subcutaneous tumor growth, but not tumor growth in the bone [31]. Here, we validated this CRPC phenotype using enzalutamide (ENZ), a direct AR inhibitor and a more potent form of ADT. PCSD1 tumors were injected intrafemorally and allowed to grow for approximately 5 weeks prior to initiating treatment. While ENZ reduced the rate of tumor growth compared to vehicle, we observed no tumor regression and tumor volume continued to increase at 30 days, indicating that this PDX tumor is inherently resistant to multiple forms of ADT (Fig. 1C). CAM-DR has been shown to cause therapeutic resistance in prostate cancer [7], and the laminin-specific integrin, α6β1 (ITGα6), is the primary integrin expressed in prostate cancer [39]. Thus, we sought to determine whether integrin signaling was enhanced following ADT. Immunohistochemical staining revealed that ITGα6 was upregulated in bone-resident PCSD1 tumors following ENZ treatment, suggesting activation of this survival pathway (Fig. 1D). BNIP3, an established target of hypoxia, was also up-regulated in the treated tumors (Fig. 1D). Next, we tested whether this model also retains the PI3K inhibitor-resistant phenotype displayed by the patient. PCSD1 tumors constitutively expressing luciferase were injected intrafemorally and allowed to establish prior to initiating treatment with 50 mg/kg SF1126, a pan-PI3K inhibitor [40], three times a week for 4 weeks. Bioluminescent imaging showed no significant change in tumor volume following treatment with vehicle or SF1126 (Fig. 1E), although the osteoblastic lesion area was reduced by PI3K inhibition (Fig. 1F). Immunohistochemical analysis of vehicle vs SF1126 treated tumors revealed a significant increase in ITGα6, BNIP3, as well as phosphorylation of Akt (S473), which indicates reactivation of the PI3K pathway. Taken together, these data demonstrate that bone-resident CRPC tumors are exposed to extensive hypoxia, display resistance to PI3K inhibition, and activate TME-driven survival signaling pathways in response to standard therapies.

### ITGα6 is activated in the bone TME and promotes resistance to PI3K inhibitors

To determine if ITGα6 is involved in conferring resistance in CRPC, ITGα6 levels in several castration-resistant LuCaP PDX tumors were compared to parental lines. While ITGα6 and ITGβ1 were present in both the parental and castration-resistant PDX models, both mRNA and protein expression were significantly higher in the castration-resistant tumors (Fig. 2A and B), suggesting the importance of this pathway for the progression to CRPC. Because the bone is known to be enriched in laminin [41, 42], we tested whether activation of ITGα6 and adhesion to laminin could account for resistance to PI3K inhibition in the bone TME. To determine the level of ITGα6 expression in bone-resident tumors vs. subcutaneous tumors, we stained tissue from the previously described cell lines and PDX models. ITGα6 levels were significantly higher in bone-resident tumors than in subcutaneous tumors (Fig. 2C and D), indicating ITGα6 is robustly expressed in the bone environment compared to soft tissue. Because we previously demonstrated that the bone TME is hypoxic, we investigated whether ITGα6 protein levels were altered in response to hypoxia. C4-2 and LNCaP cells were cultured in normoxia (20% O_2_) or hypoxia (1.0% O_2_) for 24 h and lysates were collected for immunoblotting. Hypoxia significantly increased the levels of HIF-1α and both ITGα6 and ITGβ1 (Fig. 2E), suggesting that this signaling axis is hyperactivated in the bone due to hypoxia.

**Figure 2.**
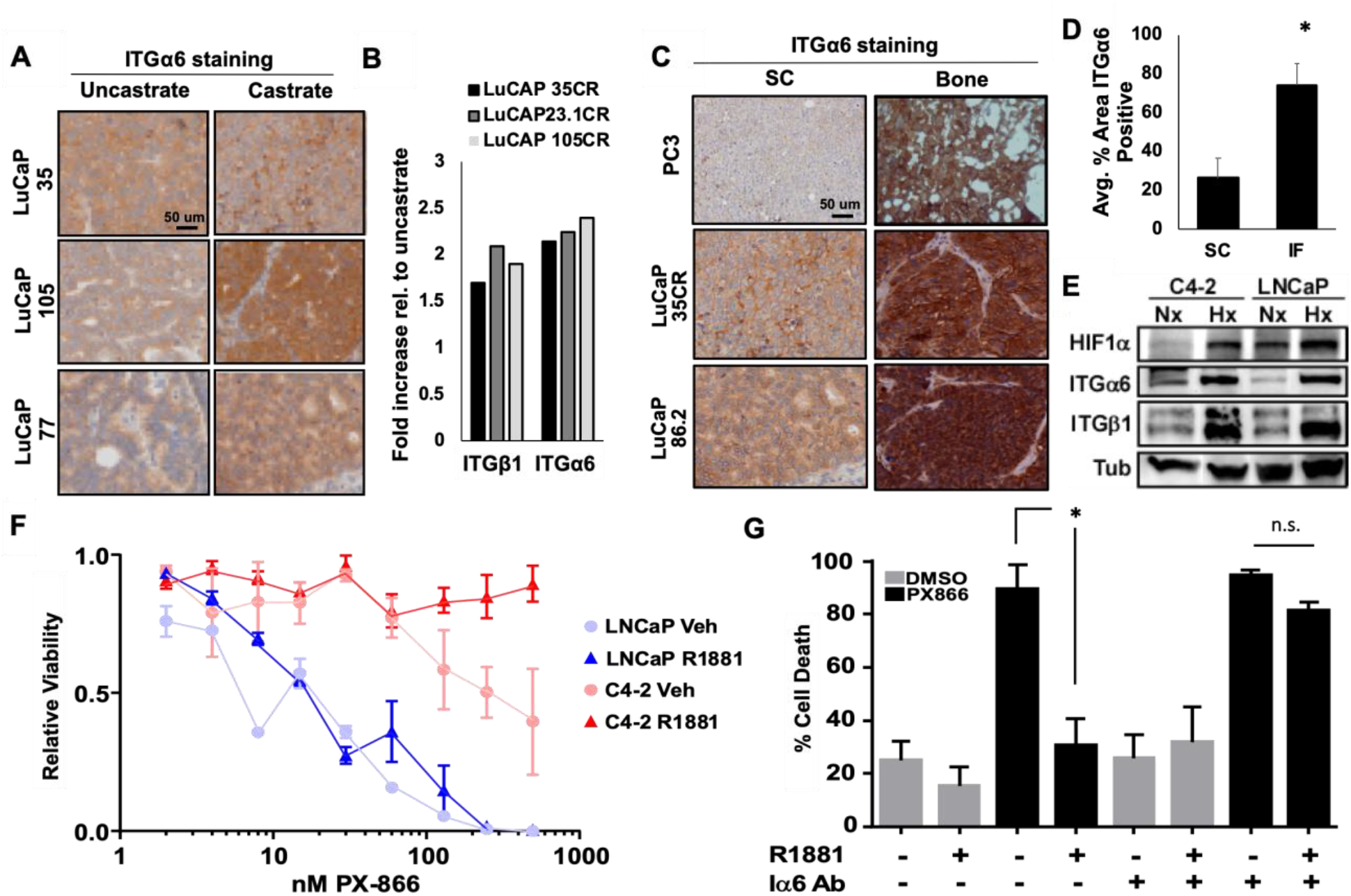
ITGα6 confers resistance to PI3K inhibitors. **A)** Immunohistochemical staining of integrin α6 (ITGα6) in castrate-resistant LuCaP (castrate) PDX tumors relative to uncastrated parental tumors. **B)** *ITGα6* and *ITGβ1* mRNA expression, from microarray data, in castration-resistant LuCaP PDX tumors relative to uncastrated parental tumors. **C)** PC3 cells or two different LuCaP PDX tumors were implanted subcutaneously (SC) or intratibially (Bone) and grown for 4 weeks. Tumors from each group were fixed and immunostained for ITGα6 and **D)** positive staining was quantified. **E)** LNCaP or C4-2 cells adherent to laminin were cultured in hypoxia for 24 h and levels of HIF-1α, ITGα6, ITGβ1, and Tubulin (Tub) protein assessed by immunoblotting. **F)** LNCaP and C4-2 cells adherent to laminin were stimulated with 10 nM R1881 for 24 h prior to and in the presence of increasing concentrations of PX866, and cell viability measured by trypan blue exclusion 48 h later. **G)** C4-2 cells were plated on laminin with or without ITGα6-blocking antibody (Iα6 Ab), then treated with or without 10 nM R1881 for 24 hours prior to and during 48 h treatment with 400 nM PX866 or DMSO (control). Cell death was assessed by trypan blue exclusion. * *p* < 0.05; n = 3; error bars indicate SEM.

Next, we tested whether integrin-mediated adhesion to laminin is sufficient to impart resistance to PI3K inhibitors. First, parental androgen-sensitive LNCaP cells and castration-resistant C4-2 cells were grown on laminin and treated with increasing concentrations of PX866 (a type I-specific PI3K inhibitor), and viability was measured after 48 h of treatment. ITGα6 is an established target of AR, so each cell line was treated with vehicle or R1881, a synthetic androgen, for 24 h to increase ITGα6 expression prior to the addition of PI3K inhibitor. Strikingly, castration-resistant C4-2 cells were 10 times more resistant to PI3K inhibition than the parental LNCaP cells (LD_50_ 20 nM vs 200 nM) and completely resistant in the presence of androgen (Fig. 2F). To determine whether AR-mediated resistance to PI3K inhibition is dependent on ITGα6, C4-2 cells were plated on laminin and serum-starved for 24 h in the presence or absence of ITGα6-blocking antibody (Iα6 Ab). Then, cells were treated with vehicle or R1881 for 24 hours prior to the addition of PX866 (400 nM) or vehicle. Treatment with PX866 in the absence of androgen induced massive cell death (greater than 80%), which was rescued by androgen stimulation (Fig. 2G). However, androgen was unable to rescue PX866-induced death in the presence of the ITGα6 inhibitor. These data indicate that ITGα6 is highly expressed in bone-resident CRPC tumors and that AR-mediated resistance to PI3K inhibition is dependent on ITGα6.

### Combined inhibition of PI3K and ITGα6 is not sufficient to completely block CRPC tumor growth

Based on our findings showing a role for ITGα6 in PI3K inhibitor resistance, we tested the efficacy of combined inhibition of ITGα6 and PI3K using *in vivo* PDX models of CRPC. First, we tested whether PTEN status altered the response of combined inhibition of PI3K and ITGα6. Mice were injected with either LuCaP 23.1 (PTEN-positive) or LuCaP 35 (PTEN-negative) PDX tumors. Tumors were allowed to grow to an average volume of 100 mm^3^ prior to randomization and treatment with an ITGα6-blocking antibody (ITGα6 Ab), PX866 (PI3K inhibitor), or the combination. In the PTEN-negative PDX tumor model, combined inhibition of both ITGα6 and PI3K was required to reduce tumor growth 2-fold, whereas inhibiting either alone had no effect (Fig. 3A). However, neither agent, alone or in combination, significantly suppressed growth of the PTEN-positive LuCaP 23.1 PDX tumors (Fig. 3B). Therefore, PTEN-negative tumors are more likely to respond to combined inhibition of ITGα6 and PI3K.

**Figure 3.**
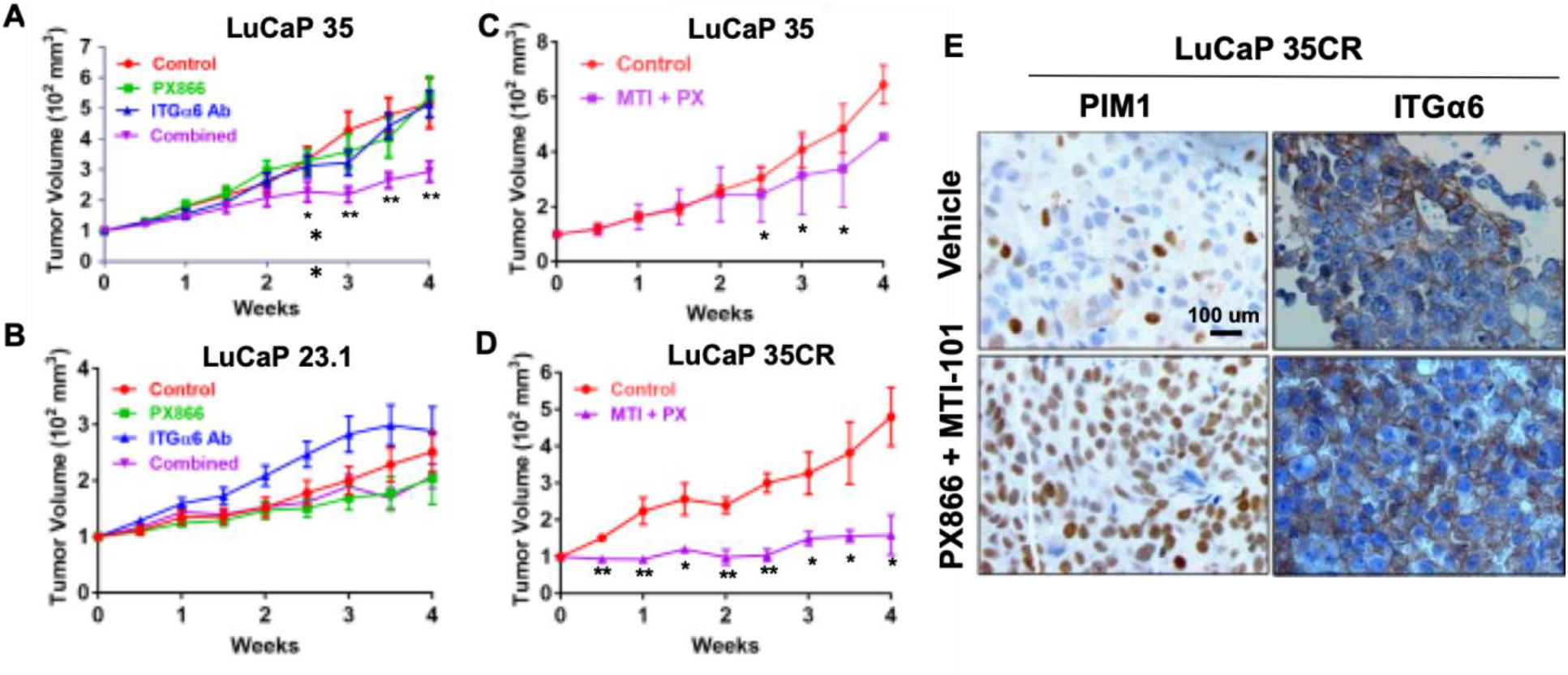
Combined inhibition of ITGα6 and PI3K reduces tumor growth but increases PIM1 expression. **A)** LuCaP 35 (PTEN-negative) and **B)** LuCaP 23.1 (PTEN-positive) human PDX tumors were grown to ~100 mm^3^ prior to treatment with vehicle (Control), 2 mg/kg PX866 (p.o. 3x/wk), 10 µg GoH3 anti-integrin α6 antibody (ITGα6 Ab) (i.p. 1x/wk), or both (Combined) for 4 wks. **C)** LuCaP 35 or **D)** LuCaP 35CR human PDX tumors were treated with vehicle (Control) or 2 mg/kg PX866 (p.o. 3x/wk) plus 40 mg/kg MTI-101 (i.p. 3x/wk) for 4 wks. In all experiments, tumor volume was measured by caliper 2x/wk. Data are normalized to body weight and initial tumor volume of 100 mm^3^. * p < 0.05, **, p < 0.005; n = 14; error bars indicate SEM. **E)** Immunostaining of PIM1 and integrin α6 (ITGα6) in tumors isolated from LuCaP 35CR treated tumors.

Recently, we developed a novel, clinically accessible ITGα6 inhibitor that shows improved efficacy in targeting CAM-DR by acting as a super agonist [43, 44]. MTI-101 is a cyclized and pharmacological peptidomimetic derived from the linear HYD1 and RZ-3 peptides [43–47], which binds integrin α6 but does not interfere with cell adhesion. To test the combined efficacy of PI3K inhibitor and MTI-101, LuCaP 35 and LuCaP 35CR PDX tumors were injected SC into mice and allowed to establish. The mice were randomly segregated into groups that were treated with either vehicle or the combination of PX866 and MTI-101. Combined treatment with PX866 and MTI-101 showed anti-tumor effects, particularly in the LuCaP 35CR PDX model, significantly reduced tumor volume over 4-fold compared to control (Fig. 3C and D). Thus, targeting ITGα6 increases the efficacy of PI3K inhibitors *in vivo*, and this strategy is effective in PTEN-negative tumors. While this drug combination significantly reduced tumor volume, it did not cause tumor regression, indicating that additional survival mechanisms are present. Immunohistochemical staining of tumor tissues from the treated mice revealed there was not much change in ITGα6 levels (Fig. 3E). Because PIM1 kinase was recently described as a mechanism of resistance to PI3K inhibitors [28, 29], we stained tumor tissues for PIM1. PIM1 was highly increased in the treated tumors (Fig. 3E). Therefore, PIM1 induction represents a novel survival pathway that could allow CRPC tumors to evade cell death following combined therapy targeting of ITGα6 and PI3K.

### Hypoxia-inducible PIM kinase promotes resistance to PI3K inhibitors

Hypoxia is known to promote drug resistance through pleotropic mechanisms. Our group recently described the importance of hypoxia-inducible PIM kinase signaling for prostate cancer cell survival in hypoxia via induction of Nrf2 [26, 48]. Because PIM levels are significantly upregulated in hypoxia, we examined PIM1 expression in subcutaneous and bone-resident PC3 and LuCaP tumor sections to determine whether PIM1 is also preferentially activated in the bone TME. As expected, because of the hypoxic nature of the bone, PIM1 expression was significantly higher in bone-resident tumors than in subcutaneous tumors (Fig. 4A and B). Next, C4-2 cells were subjected to normoxia or hypoxia (1.0% O_2_) for 6 h and lysates were collected for western blotting. As expected, hypoxia significantly increased HIF-1α, Nrf2, and all three PIM isoforms (Fig. 4C). These data indicate that hypoxia induces PIM kinase expression in CRPC, particularly in bone-metastatic tumors.

**Figure 4.**
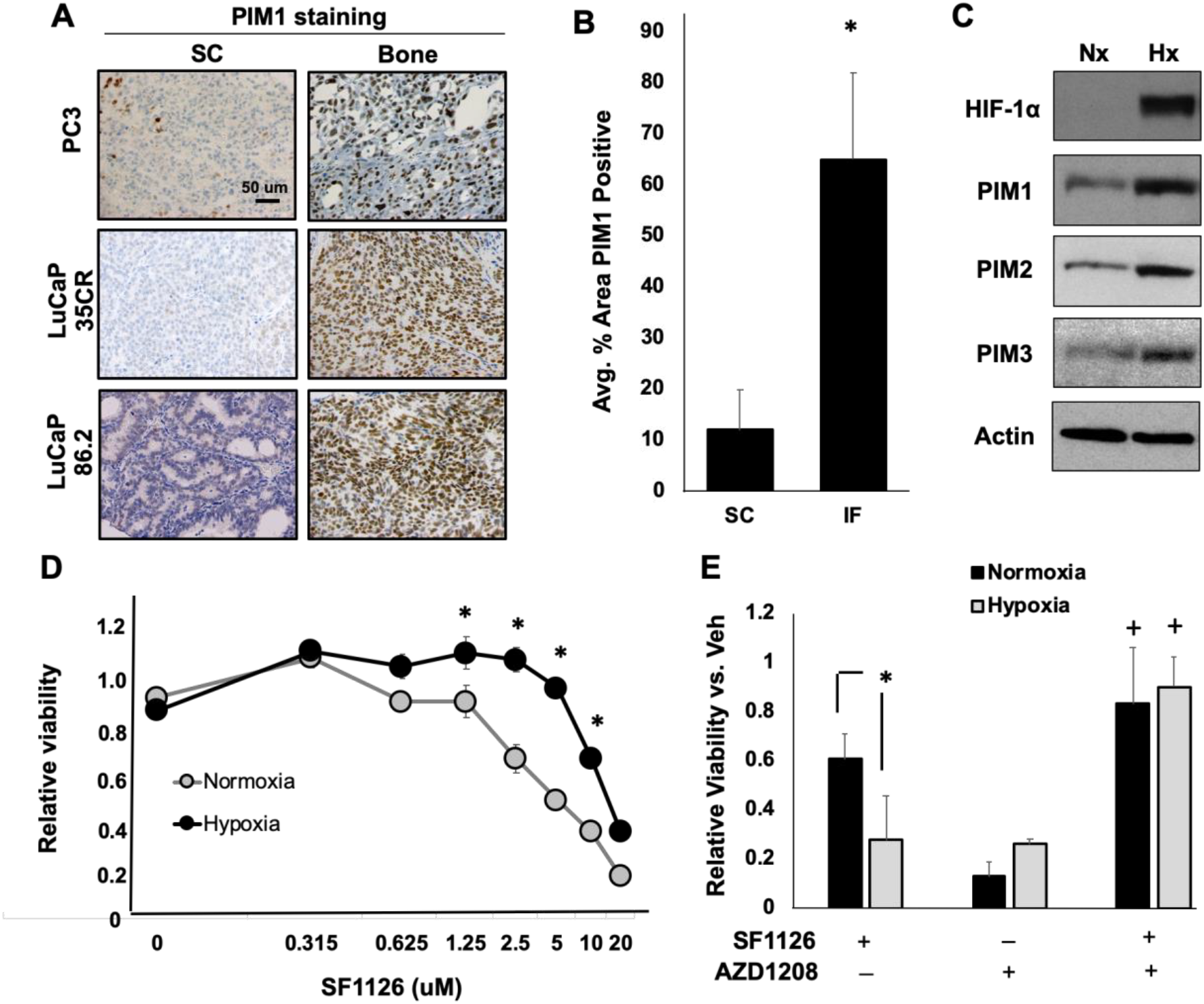
Hypoxia-inducible PIM kinase promotes resistance to PI3K inhibitors. **A)** PC3 cells or two different LuCaP PDX tumors were implanted subcutaneously (SC) or intrafemorally (IF) and grown for 4 weeks. Tumors from each group were fixed and immunostained for PIM1, and **B)** positive staining was quantified. **C)** C4-2 cells were cultured in normoxia (Nx) or hypoxia (Hx) for 24 h, and HIF-1α, Nrf2, and PIM1, 2, and 3 kinase expression assessed by immunoblotting. **D)** C4-2 cells were cultured in normoxia or hypoxia with increasing concentrations of SF1126, and cell viability quantified by crystal violet staining. **E)** C4-2 cells in normoxia or hypoxia were treated with SF1126 (5 µM), AZD1208 (1 µM), or both for 72 hours, and cell viability measured by crystal violet staining. * *p* < 0.05; n = 3; error bars indicate SEM.

To assess whether hypoxia could alter the sensitivity of CRPC cell lines to PI3K inhibitors, C4-2 CRPC cells were treated with SF1126 for 48 h in normoxia or hypoxia (1.0% O_2_), and viability was measured by crystal violet staining. Strikingly, C4-2 cells cultured in hypoxia were significantly less sensitive to SF1126 than those treated in normoxia (IC_50_ Nx = 5.3 µM; IC_50_ Hx = 12.8 µM), indicating that hypoxia can impart resistance to PI3K inhibitors in CRPC (Fig. 4D). To determine the importance of PIM activity for hypoxia-mediated resistance to PI3K inhibitors, C4-2 cells were incubated in normoxia or hypoxia for 48 h in the presence or absence of AZD1208 (a pan-PIM kinase inhibitor) alone or in combination with SF1126. Hypoxia significantly reduced cell death in response to SF1126 (Nx = 60% cell death; Hx = 28% cell death). Strikingly, AZD1208 restored the efficacy of SF1126 and increased cell death in both normoxia and hypoxia (greater than 80% cell death) (Fig. 4E). These data suggest that hypoxia-mediated resistance to PI3K inhibitors is dependent on PIM kinase activation.

### Combined inhibition of PI3K, ITGα6, and PIM kinase displays synergistic anti-cancer activity by increasing oxidative stress

To determine whether combined inhibition of PIM and ITGα6 represents an effective strategy to overcome TME-mediated resistance to PI3K inhibitors in CRPC, C4-2 cells were grown in normal conditions (normoxia on plastic) or environmental conditions that more accurately reflect the bone TME (hypoxia on laminin) and treated with SF1126, MTI-101, or AZD1208 alone or in the indicated combinations for 48 h. Under normal environmental conditions, SF1126 induced significant cell death (61% viable), whereas cells grown on laminin in hypoxia were less sensitive (93% viable), providing further evidence that a hypoxic, laminin-rich TME promotes resistance to PI3K inhibition (Fig. 5A). The doses of AZD and MTI-101 used are below their respective IC50 values. As a result, neither drug induced significant cell death alone in normoxia or bone representative conditions (Fig. 5A). Interestingly, PI3K inhibitor combined with either AZD or MTI-101 showed a modest increase in toxicity in normoxic conditions (60% and 45% viable, respectively), but cells cultured in hypoxia and on laminin maintained significant resistance to combined treatment. Strikingly, inhibition of PI3K, ITGα6, and PIM together displayed greater cytotoxicity than single agents or the dual combination treatments (less than 20% viable) in both normal and bone-like environmental conditions (Fig. 5A). Therefore, it is necessary to block both PIM and ITGα6 in order to restore sensitivity to PI3K inhibition under conditions that reflect the bone environment.

**Figure 5.**
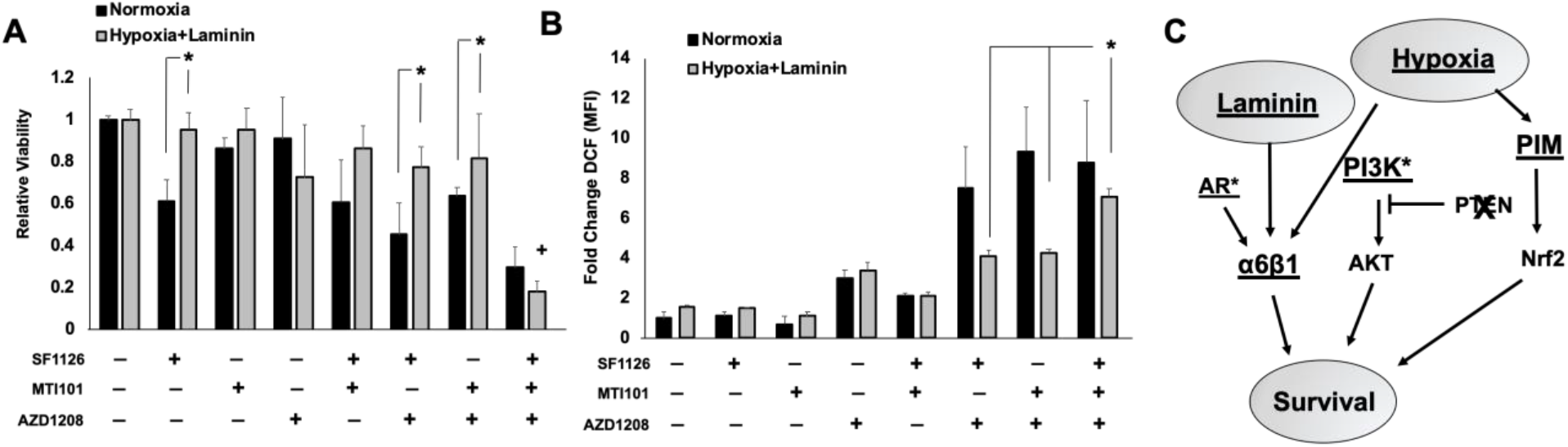
Combined inhibition of ITGα6 and PIM displays synergistic cell killing and increased oxidative stress. **A)** C4-2 cells were cultured in normoxia on plastic or in bone-like conditions, i.e. in hypoxia on laminin, and treated with SF1126 (5 µM), AZD1208 (1 µM), MTI-101 (10 µM), or all three for 48 h. Viability was assessed by crystal violet staining. **B)** C4-2 cells were treated for 24 h with the indicated drugs at the doses described in panel A prior to staining with DCF, and reactive oxygen species generation was measured by flow cytometry. **C)** Model describing the parallel signaling axes in which laminin and hypoxia in the bone tumor microenvironment activate PIM and ITGα6 to promote resistance to PI3K inhibitors in CRPC. * p < 0.05, +, p < 0.05 compared to all other treatment conditions; n = 3; error bars indicate SEM.

Both PIM kinases and ITGα6 have been implicated in reducing oxidative stress in cancer cells through independent mechanisms: PIM though activation of the antioxidant transcription factor Nrf2 and ITGα6 through the induction of autophagy [49, 50]. Therefore, we tested whether combined targeting of ITGα6 and PIM causes a significant increase in ROS under the previously described TME conditions. C4-2 cells were grown in normal or bone-like conditions and treated with the indicated drug combinations for 24 h, and DCF staining was performed to detect intercellular ROS. In vehicle-treated samples, growth on laminin and hypoxia slightly increased ROS production, and only AZD treatment was able to significantly increase ROS production as a single agent, regardless of environmental conditions (Fig. 5B). Notably, combined treatment of AZD with SF1126 or MTI-101 increased ROS dramatically in normoxia, whereas there was no additive effect in cells grown in bone-like conditions (Fig. 5B). Importantly, combined treatment with all three drugs significantly increased ROS by 7-fold compared to controls, similar to levels observed in normoxic conditions (Fig. 5B). Thus, factors enriched in the bone microenvironment prevent the accumulation of oxidative stress in CRPC cells and promote resistance to PI3K inhibition (Fig 5C). Taken together, these data provide an important preclinical rationale for the combined targeting of PIM, ITGα6, and PI3K in patients with PTEN-negative bone-metastatic CRPC.

## DISCUSSION

Treatment options for CRPC patients that have failed ADT are limited to standard chemotherapy, which is ineffective and does not markedly prolong patient survival. Surprisingly, many single-agent targeted therapies, even those whose targets are highly elevated in CRPC, have failed to improve overall survival. These findings suggest that our preclinical models do not accurately represent the patient response to therapy and that genetic alterations in tumors are not the only factors influencing sensitivity to targeted therapies, particularly in metastases. One of the key pieces of knowledge put forth by this study is that it is essential to develop and use models that mimic the bone-metastatic CRPC TME for therapeutic testing. Traditional subcutaneous xenografts and prostate xenografts do not adequately represent the bone TME. Therefore, to determine whether a drug regimen is likely to be successful for CRPC patients with bone metastasis, it is minimally essential to use bone xenograft models.

Many factors within the TME have been reported to influence prostate tumor survival and response to therapy [51]. Despite the fact that CRPC commonly metastasizes to bone, how the bone TME influences the response of prostate cancer to therapy has not been well established. In this study, we directly compared xenograft tumors from prostate cell lines and CRPC PDX models to identify survival signaling pathways that are activated by the bone TME. Investigation of these models led to the identification of environmental factors enriched in the bone, the ECM (laminin) and hypoxia, that drive resistance to PI3K inhibitors. *In vitro* modeling of “normal” and “bone-like” conditions demonstrated that the environmental characteristics of the bone TME significantly reduce the cytotoxic effect of PI3K inhibitors in CRPC models. Further investigation into the biology associated with these TME conditions led to the identification of ITGα6 and PIM kinase as key survival signaling pathways that are activated by laminin and hypoxia. Importantly, drugs that block these pathways are currently being tested in clinical trials and have proven safe and effective in humans, allowing for more rapid and straightforward clinical translation for use in CRPC patients.

Integrin α6β1 is a direct target of AR that promotes prostate cancer survival in laminin-rich TMEs such as the prostate, lymph nodes, and bone. ITGα6 has been shown to promote survival through pleotropic mechanisms, including induction of Bcl-xL and prevention of oxidative stress [39]. As seen in other models [52, 53], hypoxia also induces integrins α6 and β1 in prostate cancer. Based on these data, we propose that not only is the integrin α6β1 survival pathway active in CRPC via AR, but also under hypoxic conditions, such as those observed in the bone TME. Using the PTEN-negative LuCaP 35 PDX model, combined inhibition of both ITGα6 and PI3K was required to reduce tumor growth 2-fold *in vivo*, demonstrating the importance of both pathways in prostate cancer survival in PTEN-negative tumors. However, these agents alone or in combination did not significantly suppress growth of the PTEN-positive LuCaP 23.1 PDX tumors, suggesting that patient stratification based on PTEN expression is critical to determine who will benefit from combined PI3K therapies.

The second bone-enriched environmental condition identified in our study is hypoxia. Hypoxia conferred resistance to PI3K inhibitors in CRPC cells, and bone-resident tumors displayed significantly higher levels of HIF-1α and PIM1, which have been established as independent hypoxia-inducible proteins. PIM is particularly interesting as a therapeutic target for hypoxia, because it is upregulated in a HIF-1-independent manner. PIM is known to redundantly phosphorylate many of the pro-survival targets activated by PI3K, such as the anti-apoptotic factor Bad [54]. Thus, PIM can substitute for the survival function of PI3K, and it is essential to be aware of cellular contexts in which PIM is highly active to anticipate resistance to PI3K inhibitors. Upregulation of PIM kinase in hypoxia promotes prostate cancer cell survival in response to oxidative stress, and small molecule inhibitors of PIM have been shown to selectively kill hypoxic prostate cancer cells *in vitro* and *in vivo* by increasing ROS to toxic levels [55–58]. Importantly, inhibition of PIM in hypoxia restored the sensitivity of hypoxic CRPC cells to PI3K inhibition, further supporting the role of hypoxia-inducible PIM kinase expression in resistance to PI3K inhibitors.

In summary, we have identified novel signaling axes that are hyperactivated in the bone TME. Integrin α6β1-mediated adhesion to laminin and hypoxia-induced PIM kinase expression are parallel signaling pathways that promote survival and resistance to PI3K inhibition in CRPC. Within the bone TME, both ITGα6 and PIM1 are highly expressed and active. Importantly, our data suggest that ITGα6 and PIM kinase promote survival through independent but parallel mechanisms (Fig. 5C). PIM stimulates Nrf2 activation to block oxidative damage [55, 59], and ITGα6 counteracts oxidative stress through the induction of autophagy [50]. Therefore, we predict that each of these pathways partially compensate for the other when only one, or even two, are inhibited. This model provides an explanation for how ADT resistance develops and why inhibiting PI3K, PIM kinase, or ITGα6 alone is not effective and provides rationale for targeting multiple pathways simultaneously to improve the patient response.

## ACKNOWLEDGEMENTS

We would like to acknowledge the gift of integrin α6 antibodies AA6A and AA6ANT from Dr. Anne Cress and Dr. Scott Peterson, when at Oncothyreon, for the PX866 drug for the *in vivo* studies. We wish to thank the core services of TACMASR, EMSR, Flow Cytometry, and Genomics at the University of Arizona Cancer Center for their help and guidance. Studies were supported by funding from the NIH (CA154835 and P30CA023074), Accelerate for Success Award from the TRIF initiative at University of Arizona, and the Van Andel Research Institute.

